# A molecular pore spans the double membrane of the coronavirus replication organelle

**DOI:** 10.1101/2020.06.25.171686

**Authors:** Georg Wolff, Ronald W.A.L. Limpens, Jessika C. Zevenhoven-Dobbe, Ulrike Laugks, Shawn Zheng, Anja W. M. de Jong, Roman I. Koning, David A. Agard, Kay Grünewald, Abraham J. Koster, Eric J. Snijder, Montserrat Bárcena

**Affiliations:** Department of Cell and Chemical Biology, Section Electron Microscopy, Leiden University Medical Center, Leiden 2333 ZC, The Netherlands; Department of Medical Microbiology, Molecular Virology Laboratory, Leiden University Medical Center, Leiden 2333 ZA, The Netherlands; Department of Structural Cell Biology of Viruses, Centre for Structural Systems Biology, Heinrich Pette Institute, Leibnitz Institute of Experimental Virology, University of Hamburg, Hamburg, Germany; Department of Biochemistry and Biophysics, Howard Hughes Medical Institute, University of California San Francisco, San Francisco, CA 94143, United States; Department of Biochemistry and Biophysics, University of California San Francisco, San Francisco, CA 94143, United States; Department of Chemistry, MIN Faculty, Universität Hamburg, Hamburg, Germany

## Abstract

Coronavirus genome replication is associated with virus-induced cytosolic double-membrane vesicles, which may provide a tailored micro-environment for viral RNA synthesis in the infected cell. However, it is unclear how newly synthesized genomes and mRNAs can travel from these sealed replication compartments to the cytosol to ensure their translation and the assembly of progeny virions. Here, using cellular electron cryo-microscopy, we unveiled a molecular pore complex that spans both membranes of the double-membrane vesicle and would allow export of RNA to the cytosol. A hexameric assembly of a large viral transmembrane protein was found to form the core of the crown-shaped complex. This coronavirus-specific structure likely plays a critical role in coronavirus replication and thus constitutes a novel drug target

---

Severe acute respiratory syndrome coronavirus 2 (SARS-CoV-2) constitutes the third and most impactful example of a potentially lethal coronavirus infection in humans within the last 20 years [1-3]. The realization that such zoonotic transmissions will remain a public health threat has resurged the need to dissect coronavirus molecular biology. Coronaviruses are positive-stranded RNA (+RNA) viruses that replicate their unusually large genomes in the host cell’s cytoplasm. This process is supported by an elaborate virus-induced network of transformed endoplasmic reticulum (ER) membranes known as the viral replication organelle (RO) [4-7]. Double-membrane vesicles (DMVs) are the RO’s most abundant component and the central hubs for viral RNA synthesis [5], which includes genome replication and the synthesis of a nested set of subgenomic mRNAs. The DMV’s interior accumulates double-stranded (ds) RNA [4, 5], presumably intermediates of viral RNA synthesis, and may constitute a favorable micro-environment where factors relevant for RNA synthesis accumulate and viral dsRNA is shielded from innate immune sensors. Intriguingly, however, coronavirus-induced DMVs have been characterized as compartments that lack openings to the cytosol [4-6], despite the fact that newly-made viral mRNAs need to be transported to the cytosol for translation. Moreover, the coronavirus genome needs to be packaged by the cytosolic nucleocapsid (N) protein, before being targeted to budding sites on secretory pathway membranes, where progeny virions assemble [8].

To date, the structure of coronavirus-induced ROs in their native host cellular environment has not been analyzed by electron cryo-microscopy (cryo-EM). The murine hepatitis coronavirus (MHV) is a well-studied model for the genus *betacoronavirus*, which also includes SARS-CoV, MERS-CoV, and SARS-CoV-2. One advantage of MHV over these class-3 agents is the absence of serious biosafety constraints, enabling its use for *in situ* cryo-EM studies. To this end, we performed electron tomography (ET) on cryo-lamellae prepared by focused ion beam milling cells in the middle stage of MHV infection, when the virus-induced RO occupies large regions of the cytoplasm. The tomograms revealed abundant perinuclear DMVs having an average diameter of 257 nm (SD ± 63 nm), occasionally interconnected or connected to the ER as part of the reticulovesicular network previously described (Fig. 1, Fig. S1) [4-7]. Additionally, striking macromolecular features that previously could not be discerned in conventional EM samples became apparent (Fig. S2-4). The DMV lumen appeared to primarily contain filamentous structures that likely correspond to viral RNA (Fig. 1, Fig. S4), part of which, at least, is expected to be present as dsRNA [4, 5]. In fact, some of these filaments contained relatively long straight stretches, which is consistent with the much higher persistence length of dsRNA over ssRNA [9] (Fig. S4).

**Fig. 1.**
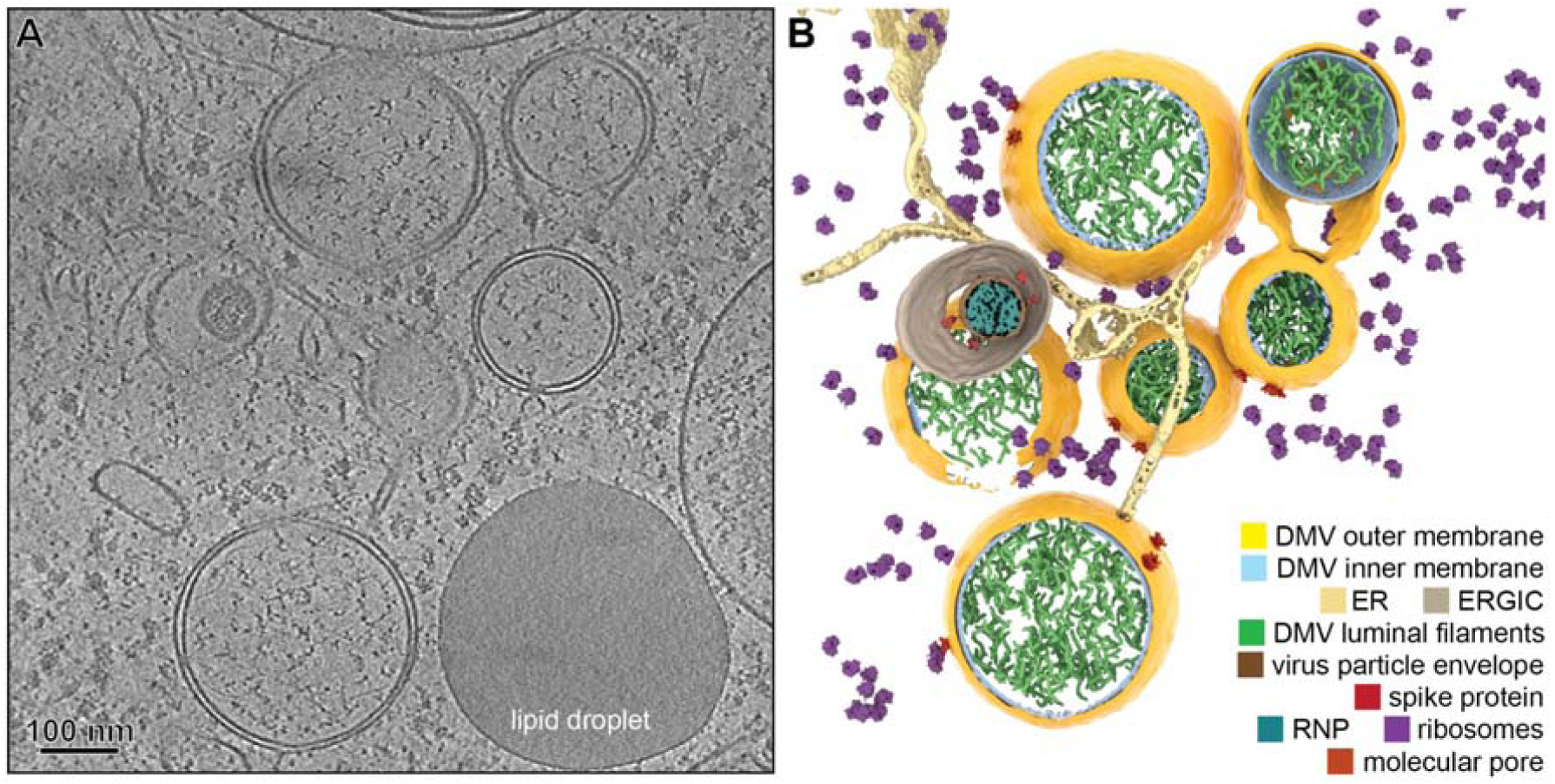
Coronavirus induced DMVs revealed by cryo-ET. (A) Tomographic slice (7 nm thick) of a cryo-lamella milled through an MHV-infected cell at a middle stage in infection. (B) 3D model of the tomogram with the segmented content annotated.

Most strikingly, our data revealed that each MHV-induced DMV contains multiple copies of a molecular complex that spans both membranes and connects the DMV interior with the cytosol (Fig. 2A). Such complexes were also found in DMVs in pre-fixed SARS-CoV-2-infected cells (Fig. 2B, Fig. S5). We surmise that this pore represents a generic coronavirus-induced molecular complex playing a pivotal role in the viral replication cycle, most likely by allowing the export of newly synthesized viral RNA from the DMV interior to the cytosol. Functionally analogous viral complexes used for RNA export include those found in the capsids of the *Reoviridae* [10] and, interestingly, the molecular pore in the neck of the invaginated replication spherules induced by flock house virus [11], although none of these is integrated in a double-membrane organelle.

**Fig. 2.**
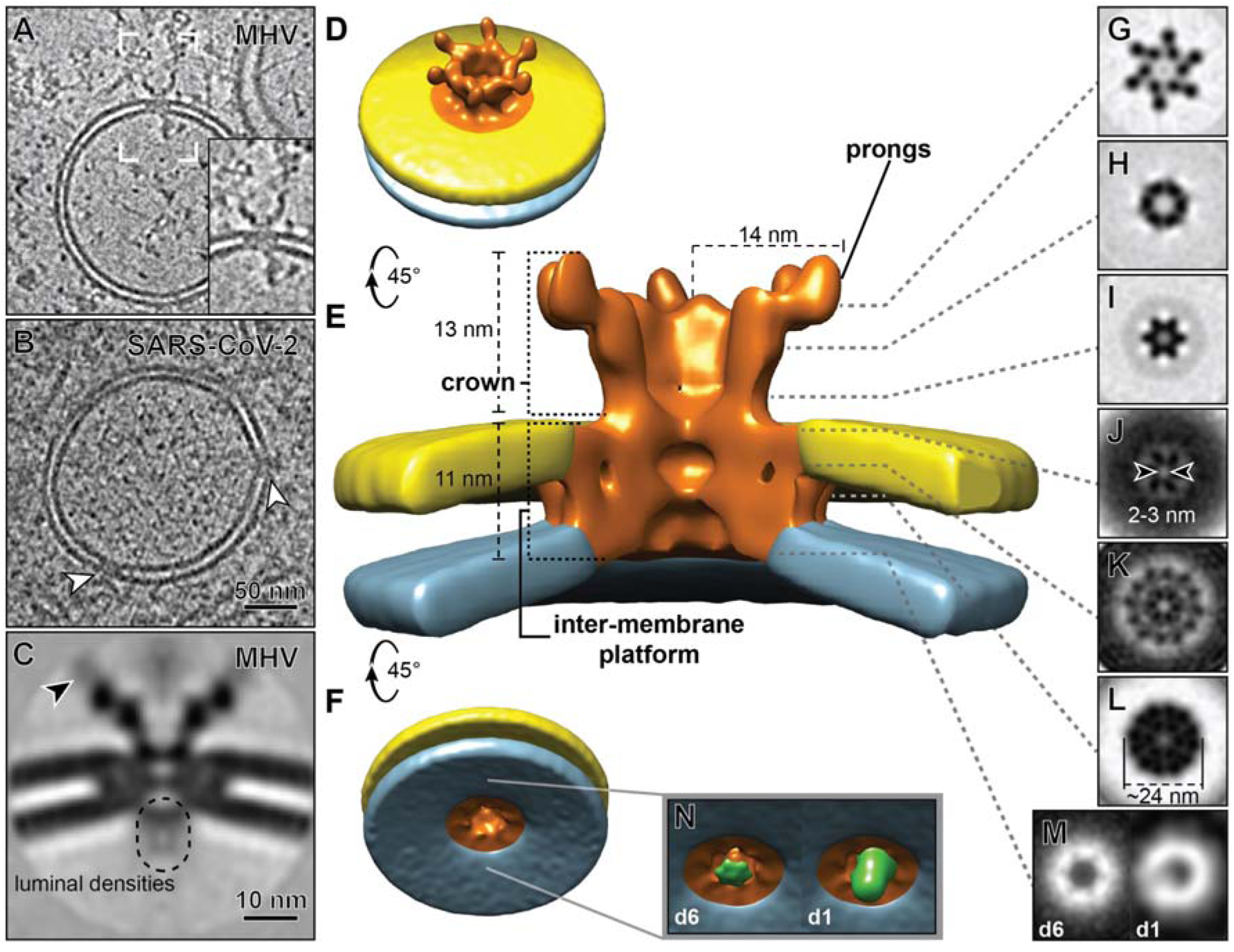
Architecture of the molecular pores embedded in DMV membranes. Tomographic slices (7 nm) revealed that pore complexes are present in both (A, inset) MHV-induced and (B) prefixed SARS-CoV-2-induced DMVs (white arrowheads). (C-L) 6-fold symmetrized subtomogram average of the pore complexes in MHV-induced DMVs. (C) Central slice through the average, suggesting the presence of flexible or variable masses near the prongs (black arrowhead) and on the DMV luminal side. (D-F) Different views of the 3D surface-rendered model of the pore complex (copper-colored) embedded in the outer (yellow) and inner (blue) DMV membranes. (G-M) 2D cross-section slices along the pore complex at different heights. (M, N) An additional density at the bottom of the 6-fold symmetrized volume (d6, green) appears as an off-centered asymmetric density in an unsymmetrized average (d1).

Subtomogram averaging of the double-membrane-spanning complexes in MHV-induced DMVs unveiled that they have an overall 6-fold symmetry (Fig. 2, Fig. S6). A cytosolic crown-like structure extends ∼13 nm into the cytosol and is based on a ∼24-nm wide platform embedded between inner and outer DMV membranes, which do not fuse and maintain a distance of ∼4.5 nm, which was the typical inter-membrane spacing found in DMVs (Fig. S2). The complex forms a channel that follows its central 6-fold axis. On the DMV luminal side, the channel starts with a ∼6-nm wide opening that narrows towards the cytosol and has two tight transition points (Fig. 2J and Fig. 2L). The one at the level of the DMV outer membrane (Fig. 2J) is the most constricted, with an opening of ∼2-3 nm, but would still allow the transition of RNA strands. Towards the cytosolic space, the complex opens up into the crown-like structure, exposing six cytosolic “prongs” that extend to a radius of ∼14 nm. With an achieved resolution of 3.1 nm (Fig. S6A), we estimate that the complex has a total molecular mass of 3 MDa, of which the crown would represent ∼1.2 MDa and each individual prong ∼30 kDa (Fig. S6C).

We then considered the possible constituents of this complex. Coronavirus genome expression starts with translation of the 5’-proximal two-thirds of the +RNA genome into two large replicase polyproteins that are proteolytically cleaved into 16 nonstructural proteins (nsps) [12]. Three of these nsps are transmembrane proteins and thus potential candidates to be components of the pore: nsp3 (222 kDa in MHV), nsp4 (56 kDa), and nsp6 (33 kDa), containing two, four and six TMDs, respectively [13-15] (Fig. 3A). These nsps are known to engage in diverse homotypic and heterotypic interactions [16] that are thought to drive the formation of double-membrane RO structures [17-19]. Based on its size, the multidomain MHV nsp3 subunit (∼2,000 amino acids) is an attractive putative constituent of the pore. MHV nsp3 consists of a large cytosolic region of ∼160 kDa followed by two TMDs and a C-terminal cytosolic domain of ∼41 kDa [13]. Whereas the TMDs and C-terminal domain are highly conserved, the domain composition and size of the N-terminal part of nsp3 is quite variable among coronaviruses, also due to the duplication of several domains in specific lineages [16, 20]. Interestingly, several nsp3 domains may interact with RNA [16], including the conserved N-terminal ubiquitin-like domain 1 (Ubl1) that binds both single-stranded (ss) RNA [21] and the N protein [22, 23].

**Fig 3.**
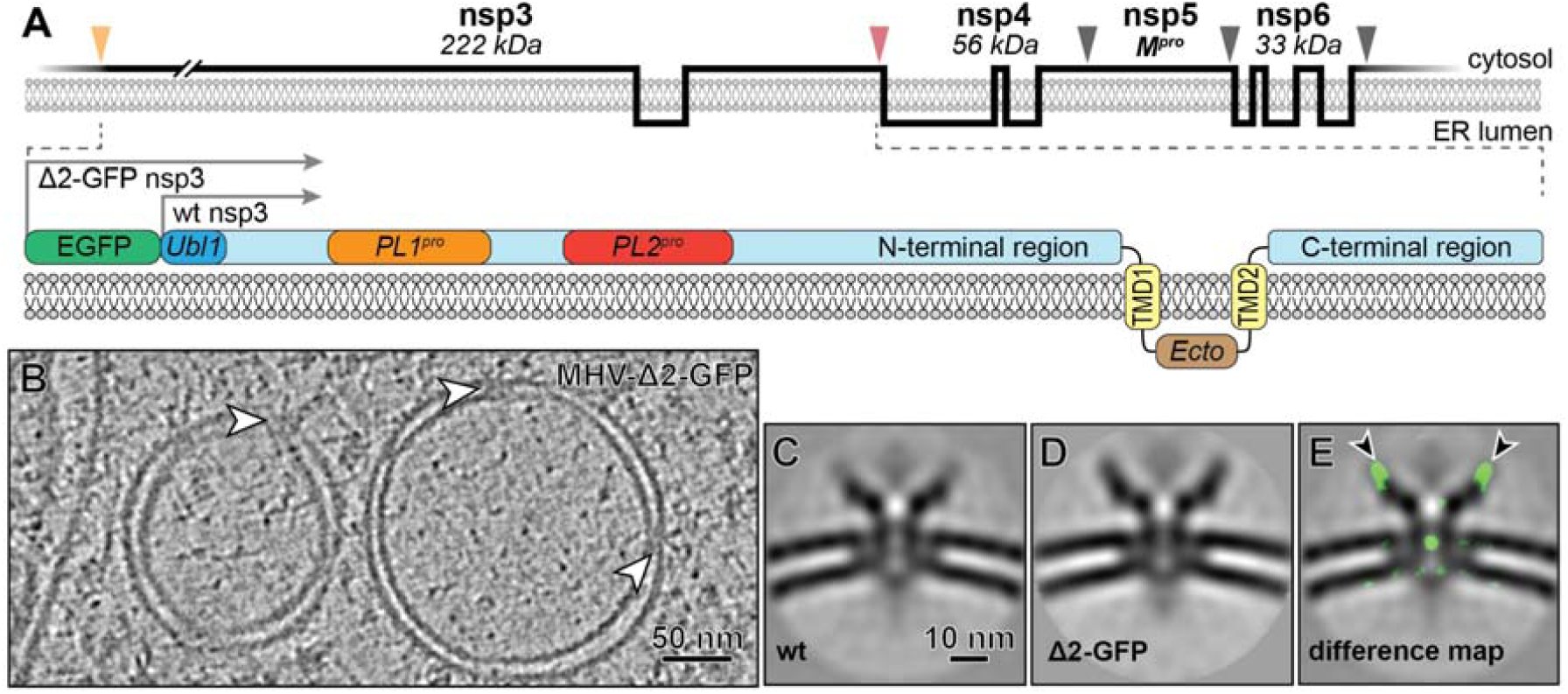
The coronavirus transmembrane-protein nsp3 is a component of the pore complex. (A) Membrane topology (top) of MHV transmembrane nsps with protease cleavage sites indicated by orange (PL1^pro^), red (PL2^pro^) and grey (M^pro^) arrowheads. (bottom) Detailed depiction of nsp3, showing some of its sub-domains and the position of the additional GFP moiety that is present in MHV-Δ2-GFP. (B) Tomographic slice of DMVs induced by MHV-Δ2-GFP with embedded pore complexes (white arrowheads). Comparison of the central slices of the 6-fold symmetrized subtomogram averages of the pore complexes in DMVs induced by (C) wt MHV and (D) MHV-Δ2-GFP. (E) Density differences of 3 standard deviations between the mutant and the wt structures, shown as a green overlay over the latter, reveal the presence of additional (EGFP) masses in the mutant complex (black arrowheads). PL^pro^, papain-like protease; M^pro^, main protease.

To investigate whether nsp3 is a component of the DMV molecular pore, we imaged cells infected with an engineered MHV mutant lacking nsp2 and expressing a nsp3 with an enhanced GFP (EGFP) moiety fused to the Ubl1 domain (MHV-Δ2-GFP [24]) (Fig. S7). Subtomogram averaging of the pore complexes in these samples (Fig. 3B) revealed the presence of additional densities on top of the prongs, representing a mass compatible with that of EGFP (Fig. 3C-E). These results unambiguously identified nsp3 as a major constituent of the complex and provided a first clue on its orientation, with the 12.6-kDa Ubl1 domain residing in the prongs of the structure. Given the observed 6-fold symmetry of the complex, six copies of nsp3 can be envisioned to constitute most of the cytosolic crown-like structure (∼1.2 MDa). Likely, other viral and/or host proteins are also involved in the formation of the ∼1.8 MDa inter-membrane platform, with the nsp4 and nsp6 transmembrane subunits being prominent candidates. In fact, studies in expression systems suggest that nsp3-nsp4 interactions could drive the membrane pairing that may be the initial step in DMV formation [17-19], while mutagenesis of the luminal domain of nsp4 results in aberrant DMV formation during infection [25].

The molecular pores frequently appear to interact with other macromolecules on both the cytosolic and the DMV luminal side (Fig. S8). These interactions seem to be dynamic as in the averages they show as largely blurred out densities that appear to extend either from the cytosolic prongs or from the opening at the DMV luminal side of the pore complex (Fig. 2C). A smaller region of the latter, however, showed relatively higher density, and was resolved in unsymmetrized averages as a closely associated and slightly off-centered mass (Fig. 2M, N, Fig. S6D), which can be speculated to be part of the viral replication machinery. The replication/transcription complex (RTC) of coronaviruses is thought to consist of a subset of relatively small (∼10-110 kDa) nsps with the RNA-dependent RNA polymerase (nsp12) at its core [26-28]. However, some of these subunits may associate with the RTC only transiently and the nsp stoichiometries of the complex are unknown. The tomographic data did not reveal clear macromolecular features within the DMV lumen that would suggest the formation of larger RTC assemblies. Interestingly, however, the luminal interaction partners of the pore complex, prominent as masses varying in shape and size, appear to interact with the putative RNA content of the DMV lumen (Fig. S8).

The interaction partners of the cytosolic nsp3 prong do not show a uniform picture either, ranging from chain-like masses protruding from the prongs to larger assemblies (Fig S8, black arrowheads). The subdomains of the long N-terminal nsp3 domain engage in a range of viral and virus-host interactions [16, 20]. Consequently, the list of possible prong interactors is long, but among them the viral N protein (55 kDa) is a prominent candidate as it was shown to bind the Ubl1 domain that resides in the nsp3 prong [22, 23]. The Ubl1-N interaction was proposed to target viral RNA to replication sites early in infection [23], but it is tempting to speculate that it may also modulate RNA exit and encapsidation on the cytosolic side of the pore complex). Notably, DMV-rich regions of the cytosol were often crowded with protein assemblies with an approximate diameter of 15 nm (Fig. S9). These strongly resembled the condensed nucleocapsid structure observed in coronavirus particles, a helical ribonucleoprotein complex (RNP) consisting of the RNA genome in tight association with N protein oligomers [29] (Fig. S9).

Our findings thus suggest a pathway for viral genomic RNA from the putative site of RNA synthesis inside DMVs, via the channel of the pore into the cytoplasm, where it would be encapsidated. In our model, either during or after viral RNA synthesis, specific replicase subunits may associate with the pore complex to guide the newly synthesized RNA towards it (Fig. 4A). As proposed for other +RNA viral ROs [11], only positive-stranded RNAs would need to be exported, whereas their low abundance negative-stranded templates could remain inside the DMVs. Subsequently, on the cytosolic side, it remains an open question whether all exported viral mRNAs would associate with the N protein (Fig. 4B). Alternatively, the interaction of viral mRNAs with the accumulating N protein could serve to select part of the newly made genomes for packaging, while the remainder would be used for translation, together with the much smaller subgenomic mRNAs that constitute ∼95% of the viral RNA products [30]. Genome-containing RNPs would travel to membrane sites where the viral envelope proteins accumulate and engage in the formation of progeny virions (Fig. 4C) [31]. These bud into single-membrane compartments (Fig. 4D), typically derived from the ER-to-Golgi intermediate compartment (ERGIC) [8], and travel along the secretory pathway to be released into extracellular space.

**Fig. 4.**
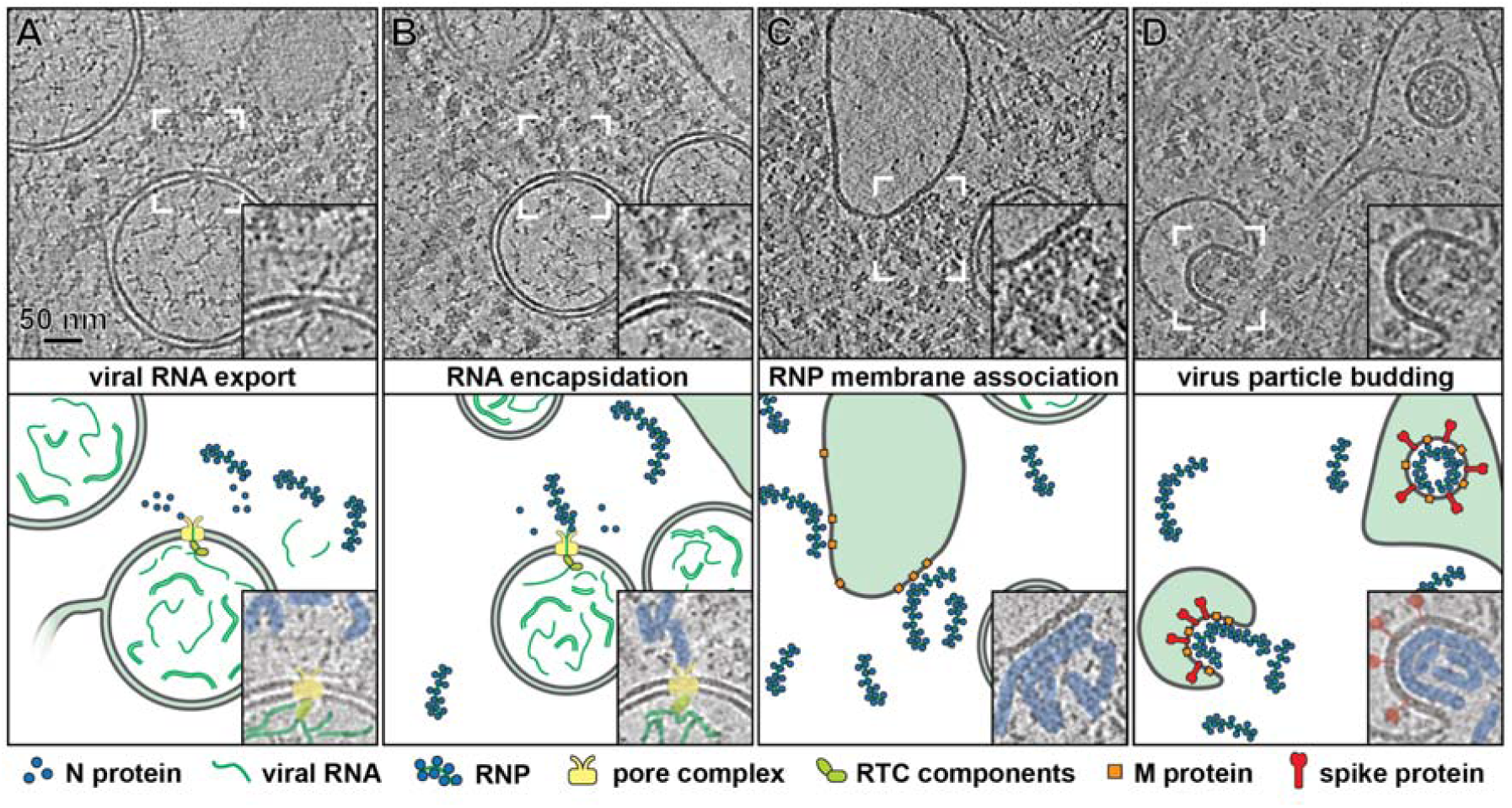
Model of the coronavirus genomic RNA transit from the DMV lumen to virus budding sites. (Top) Tomographic slices (7 nm) from MHV-infected cells highlighting the respective steps in the model (bottom). (A) The molecular pore exports viral RNA into the cytosol, (B) where it can be encapsidated by N protein. (C) Cytosolic RNP then can travel to virus assembly sites for membrane association and (D) subsequent budding of virions.

The double-membrane-spanning molecular pore revealed in this study is a unique and complex structure that could only be unveiled in its native functional context by cellular cryo-EM. This complex likely constitutes the long-sought exit pathway for coronaviral RNA products from the DMV’s interior towards the cytosol, with the large and multifunctional nsp3 being its central component. Although the exact mode of function of this molecular pore remains to be elucidated, it would clearly represent a key structure in the viral replication cycle that is likely conserved among coronaviruses, and thus may constitute a general coronavirus-specific drug target.

## Acknowledgments

We are grateful to Dr. Mark Denison for sharing MHV-Δ2-GFP and Dr. Leon Caly and Dr. Julian Druce for sharing SARS-CoV-2 isolate betaCoV/Australia/VIC01/2020. We thank Natacha Ogando for helping with the BSL3 laboratory work, Frank Faas for technical support and Stuart Howes and Thomas Sharp for advice in image processing and helpful discussions. We thank Jürgen Plitzko and Miroslava Schaffer for introducing us to cryo-FIB-milling. EM tomography data of MHV-infected samples was collected at The Netherlands Centre for Electron Nanoscopy (NeCEN) with assistance from Christoph Diebolder and Rebecca Dillard.

## Author contributions

Conceptualization: M.B., A.J.K., E.J.S.; Software: D.A.A., S.Z.; Validation: M.B., G.W.; Formal analysis: M.B.; G.W.; Investigation: M.B.; A.W.M.J.; U.L.;R.W.A.L.L.; J.C.Z.; Resources: K.G.; A.J.K.; E.J.S.; Data Curation: M.B., R.I.K., G.W.; Writing – original draft preparation: M.B., E.J.S., G.W.; Writing – review and editing: D.A.A., M.B., K.G., A.J.K., E.J.S, G.W.; Visualization: M.B., R.I.K., R.W.A.L.L., G.W.; Supervision: M.B., A.J.K., E.J.S.; Project administration: M.B.; Funding acquisition: A.J.K., E.J.S.

## References

1. Ksiazek, T.G., et al., A novel coronavirus associated with severe acute respiratory syndrome. N Engl J Med, 2003. 348(20): p. 1953–66.

2. Zaki, A.M., et al., Isolation of a novel coronavirus from a man with pneumonia in Saudi Arabia. N Engl J Med, 2012. 367(19): p. 1814–20.

3. Zhu, N., et al., A Novel Coronavirus from Patients with Pneumonia in China, 2019. N Engl J Med, 2020. 382(8): p. 727–733.

4. Knoops, K., et al., SARS-coronavirus replication is supported by a reticulovesicular network of modified endoplasmic reticulum. PLoS Biol, 2008. 6(9): p. e226.

5. Snijder, E.J., et al., A unifying structural and functional model of the coronavirus replication organelle: Tracking down RNA synthesis. PLoS Biol, 2020. 18(6): p. e3000715.

6. Maier, H.J., et al., Infectious bronchitis virus generates spherules from zippered endoplasmic reticulum membranes. MBio, 2013. 4(5): p. e00801–13.

7. Ulasli, M., et al., Qualitative and quantitative ultrastructural analysis of the membrane rearrangements induced by coronavirus. Cell Microbiol, 2010. 12(6): p. 844–61.

8. de Haan, C.A. and P.J. Rottier, Molecular interactions in the assembly of coronaviruses. Adv Virus Res, 2005. 64: p. 165–230.

9. Abels, J.A., et al., Single-molecule measurements of the persistence length of double-stranded RNA. Biophys J, 2005. 88(4): p. 2737–44.

10. Ding, K., et al., In situ structures of rotavirus polymerase in action and mechanism of mRNA transcription and release. Nat Commun, 2019. 10(1): p. 2216.

11. Ertel, K.J., et al., Cryo-electron tomography reveals novel features of a viral RNA replication compartment. Elife, 2017. 6.

12. Snijder, E.J., E. Decroly, and J. Ziebuhr, The Nonstructural Proteins Directing Coronavirus RNA Synthesis and Processing. Adv Virus Res, 2016. 96: p. 59–126.

13. Oostra, M., et al., Topology and membrane anchoring of the coronavirus replication complex: not all hydrophobic domains of nsp3 and nsp6 are membrane spanning. J Virol, 2008. 82(24): p. 12392–405.

14. Oostra, M., et al., Localization and membrane topology of coronavirus nonstructural protein 4: involvement of the early secretory pathway in replication. J Virol, 2007. 81(22): p. 12323–36.

15. Kanjanahaluethai, A., et al., Membrane topology of murine coronavirus replicase nonstructural protein 3. Virology, 2007. 361(2): p. 391–401.

16. Neuman, B.W., Bioinformatics and functional analyses of coronavirus nonstructural proteins involved in the formation of replicative organelles. Antiviral Research, 2016. 135: p. 97–107.

17. Angelini, M.M., et al., Severe Acute Respiratory Syndrome Coronavirus Nonstructural Proteins 3, 4, and 6 Induce Double-Membrane Vesicles. mBio, 2013. 4(4): p. e00524–13.

18. Oudshoorn, D., et al., Expression and Cleavage of Middle East Respiratory Syndrome Coronavirus nsp3-4 Polyprotein Induce the Formation of Double-Membrane Vesicles That Mimic Those Associated with Coronaviral RNA Replication. mBio, 2017. 8(6): p. e01658–17

19. Hagemeijer, M.C., et al., Membrane rearrangements mediated by coronavirus nonstructural proteins 3 and 4. Virology, 2014. 458-459: p. 125–35.

20. Lei, J., Y. Kusov, and R. Hilgenfeld, Nsp3 of coronaviruses: Structures and functions of a large multi-domain protein. Antiviral Research, 2018. 149: p. 58–74.

21. Serrano, P., et al., Nuclear magnetic resonance structure of the N-terminal domain of nonstructural protein 3 from the severe acute respiratory syndrome coronavirus. J Virol, 2007. 81(21): p. 12049–60.

22. Hurst, K.R., C.A. Koetzner, and P.S. Masters, Characterization of a critical interaction between the coronavirus nucleocapsid protein and nonstructural protein 3 of the viral replicase-transcriptase complex. J Virol, 2013. 87(16): p. 9159–72.

23. Cong, Y., et al., Nucleocapsid Protein Recruitment to Replication-Transcription Complexes Plays a Crucial Role in Coronaviral Life Cycle. J Virol, 2020. 94(4).

24. Freeman, M.C., et al., Coronavirus replicase-reporter fusions provide quantitative analysis of replication and replication complex formation. J Virol, 2014. 88(10): p. 5319–27.

25. Gadlage, M.J., et al., Murine Hepatitis Virus Nonstructural Protein 4 Regulates Virus-Induced Membrane Modifications and Replication Complex Function. Journal of Virology, 2010. 84(1): p. 280.

26. Kirchdoerfer, R.N. and A.B. Ward, Structure of the SARS-CoV nsp12 polymerase bound to nsp7 and nsp8 co-factors. Nature Communications, 2019. 10(1): p. 2342.

27. Sevajol, M., et al., Insights into RNA synthesis, capping, and proofreading mechanisms of SARS-coronavirus. Virus Res, 2014. 194: p. 90–9.

28. Subissi, L., et al., SARS-CoV ORF1b-encoded nonstructural proteins 12-16: replicative enzymes as antiviral targets. Antiviral Res, 2014. 101: p. 122–30.

29. Gui, M., et al., Electron microscopy studies of the coronavirus ribonucleoprotein complex. Protein Cell, 2017. 8(3): p. 219–224.

30. Ogando, N.S., et al., SARS-coronavirus-2 replication in Vero E6 cells: replication kinetics, rapid adaptation and cytopathology. 2020: p. 2020.04.20.049924.

31. Kuo, L., C.A. Koetzner, and P.S. Masters, A key role for the carboxy-terminal tail of the murine coronavirus nucleocapsid protein in coordination of genome packaging. Virology, 2016. 494: p. 100–7.

