## Supplementary Materials for "A molecular pore spans the double membrane of the coronavirus replication organelle"

### Supplementary Text:

#### Number of molecular pores per DMV

To estimate the average number of molecular pores per DMV, 8 tomograms out of 6 lamellae were used to count the number of pore complexes present in a total of 30 DMVs. Pores could be found in all the DMVs in the tomograms. The average number of detected pores (4.8 per DMV) was then extrapolated to account for the fact that only part of an average-sized DMV (257 nm in diameter, Fig. S1) is embedded in the thin cryo-lamella used for this quantification (~150 nm thick on average). This resulted in an estimate of ~8 pores per DMV, which can be considered a lower limit to the real number, as it does not take into consideration the missing-wedge effect intrinsic to electron tomography data that hampers the detection of molecular pores in certain orientations of the reconstructed volumes [17].

### Materials and Methods:

#### Cell culture, infection and vitrification

17clone1 and Vero E6 cells were grown as described previously [1]. R1/4 gold Quantifoil grids (200 mesh, Quantifoil Micro Tools, Jena, Germany) were cleaned overnight by placing them on a chloroform soaked filter paper in a sealed glass dish. After letting them dry, the grids were glow-discharged and placed in 3.5 cm diameter cell culture dishes (Greiner Bio-One, Frickenhausen, Germany) containing PBS. After UV-sterilization for 30 min in a laminar-flow, PBS was removed and 110,000 17clone1 or Vero E6 cells were seeded per dish and incubated overnight at 37 °C . 17clone1 cells were infected with wild type (wt) MHV (ATTC VR-764) or MHV-Δ2-GFP [2]. In the MHV-Δ2-GFP mutant (generously provided by Mark Denison, Vanderbilt University Medical Centre, Nashville, TN, USA), the nsp2-coding region, which is dispensable for virus replication in cell culture [3], is replaced by the EGFP-coding sequence, yielding a GFP-nsp3 fusion protein [2]. Samples were plunge-frozen in liquid ethane at 8 h post-infection (p.i.) (wt MHV) or 10 h p.i. (MHV-Δ2-GFP) with a Leica EM GP (Leica microsystems, Wetzlar, Germany) automated plunge-freezer, after applying 15 s of blotting from the backside of the grid and 1s post-blotting time in conditions of 37 °C and 95% humidity. VeroE6 cells were infected with SARS-CoV-2 isolate BetaCoV/Australia/VIC01/2020 (GeneBank ID: MT007544.1; kindly provided by Leon Caly and Julian Druce, Doherty Institute, Melbourne, Australia [4]) and chemically fixed at 8 h p.i. with 3% PFA in cell culture medium. After 1 h the original fixative was replaced by 0.5% PFA in cell culture medium and the samples stored overnight at 4 °C, before exporting the samples from the biosafety level 3 facility of Leiden University Medical Center, the

Netherlands, and then by PFA-free medium 30 min before plunge-freezing. The grids were plunge-frozen in liquid ethane using a Leica EM GP with 10 s blotting, 2 s post-blotting time and chamber conditions of 21 °C and 90% humidity and stored in liquid nitrogen until further use.

**Electron cryo-tomography**

Plunge-frozen samples were transferred to an Aquilos cryo-focused ion beam scanning electron microscope (FIB-SEM) (Thermo Fisher Scientific, Hillsboro, OR, USA) and cryo-lamellae were prepared as described [5]. For the MHV-Δ2-GFP samples, prior to FIB-milling cryo-fluorescence light microscopy (cryo-FLM) was performed on an upright FLM-system (Axioimager Zeiss, Oberkochen, Germany) equipped with a cryo-correlative light and EM stage (CMS196, Linkam, Tadworth, UK). The resulting images were integrated in the FIB-milling workflow using MAPS software (Thermo Fisher Scientific) to guide the finding of suitable milling regions (Fig. S8). A total of approximately 200 cryo-lamellae of 120-200 nm thickness were prepared (139 wt MHV, 32 MHV-Δ2-GFP, 23 SARS-CoV-2), of which about 40% contained the features of interest and were also of a sufficient quality to allow tomographic data collection. Tilt-series acquisition was carried out on FEI Titan Krios (Thermo Fisher Scientific) electron microscopes operated at 300 kV and equipped with a Gatan GIF Quantum energy filter (Gatan, Pleasanton, CA). Movie-frames were collected using a Gatan K2 (MHV infections) or Gatan K3 (SARS-CoV-2 infections) Summit direct detection device operated in counting mode. Zero-loss imaging with a 20 eV slit width was done with a pixel size of 3.51 Å or 3.37 Å (MHV and SARS-CoV-2 data, respectively) and defocus values ranging from -5 to -8 µm. Bi-directional or dose-symmetric [6] tilt-series were collected with FEI Tomo4 and serialEM software, respectively. The total dose applied on the sample was 120-140 $e^{-}/Å^{2}$, which was distributed over tilt-series covering a 100˚-120˚ range with 2˚ or 3˚ tilt increments. Recorded movie frames were aligned with MotionCor2 using 5 x 5 patches [7]. The resulting fiducialless tilt-series were aligned and SART (Simultaneous Algebraic Reconstruction Technique) reconstructed using 9 SART iterations with AreTomo (Zheng and Agard, https://msg.ucsf.edu/software). CTF-estimation was done with CTFFIND4 [8], while CTF-correction and denoising for visualization and modelling purposes was done with the ctfphaseflip [9] and the nonlinear anisotropic diffusion [10] packages included in IMOD [11], respectively.

**Subtomogram averaging**

Individual DMV-embedded molecular channels were manually picked from 4-times binned tomograms using the IMODs 3dmod interface. The coordinates were then transferred into Dynamo [12], which was used for particle cropping, alignment and averaging from 2-times binned tomograms (7.02 Å per pixel). For the wt MHV sample, 611 particles (96 px box size) were extracted out of the 53 highest quality tomograms (from 33 lamellae), while for the MHV-Δ2-GFP sample 154 particles out of 31 tomograms were cropped (from 16 lamellae). The initial alignment allowed full rotational search in all directions and resulted in a rough average that was manually centered onto the central z-axis to establish the shift to be applied to the individual particles. This was followed by a fine centering step using the 20-fold symmetrized average as a reference to fine align the particles on the central axis. Subsequent alignment steps allowed only azimuthal rotations and limited lateral shifts. For the wt MHV sample, three subsets of random particles were used to generate averages that were used for multi-reference alignment, which resulted in a class of 346 particles that provided the highest resolution and were included in the final averages. The discarded particles likely represent different conformational stages of the complex that, nevertheless, appeared to be not sufficiently populated to provide further insight at this stage. Two parallel alignments were carried out with the selected particles, with or without applying 6-fold symmetry. For all particles a tight missing wedge mask was applied that removed all information beyond ± 40˚ in Fourier space. The Fourier shell correlation curves were determined from individually refined half-datasets of the wt MHV and MHV-Δ2-GFP subtomogram averages, respectively. The correct handedness of the averages was assessed by comparison with an average of aligned ribosome particles originating from our data to a published ribosome structure [13].

**3d modelling and model interpretation**

3D surface models of the DMVs in the cellular environment were generated in Amira (Thermo Fisher Scientific) and animations in Cinema4D (Maxon Computer, Friedrichsdorf, Germany) software. Model files of ribosomes and channel complexes were generated in Dynamo, in the case of the molecular pores, from particle maps generated during sub-tomogram averaging, and in the case of ribosomes by template matching with a ribosome average generated from our data.

Surface models of the channel complex were generated in UCSF Chimera software [14] from 3d averages low-pass filtered to the resolution achieved by sub-tomogram averaging. Molecular weight of individual parts of the channel complex were estimated by calculating the (sub)volumes enclosed by the surface models generated in Chimera and dividing them by the average density of globular proteins [15]. For the generation of difference maps of the averages from the molecular channels found in wt MHV infected and MHV-Δ2-GFP infected cells, the radial power spectra of the reconstructed volumes were adjusted to each other before subtracting the volumes using the relion_image_handler function in Relion 3.1 [16].

### Supplementary Figures

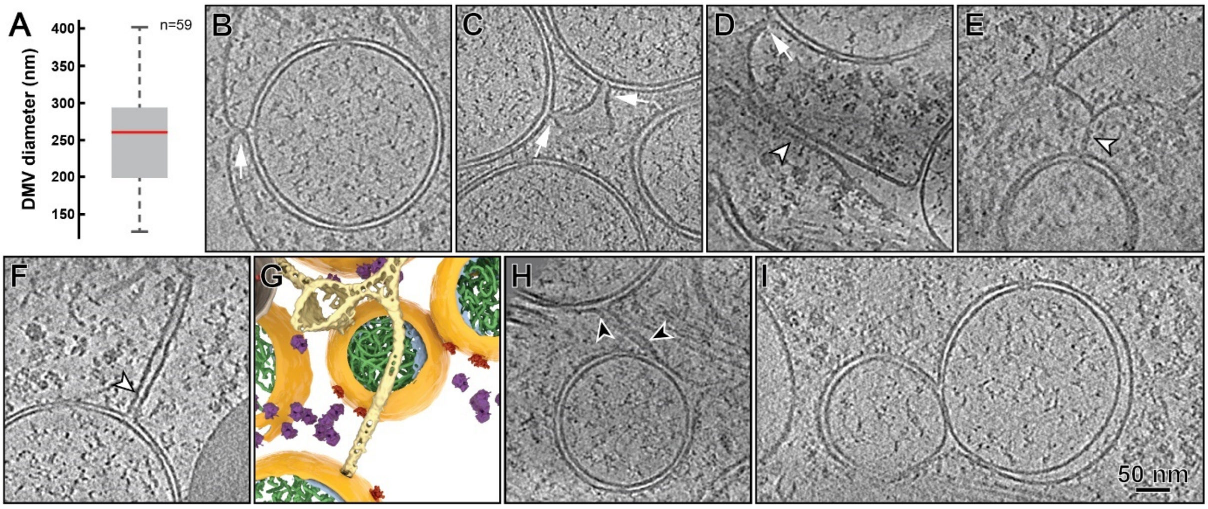


**Fig. S1.** Overall morphology of the MHV-induced DMVs in cryo-lamellae. (A) Whisker plot of the size distribution of the DMVs as measured from the tomograms at the DMV equators. Most of the DMVs in the cryo-EM samples were clearly spherical, a subset of which was used for the plot. (B-G) Connections between the outer membrane of the DMVs and the ER were occasionally detected in the tomograms. These include the neck-like connections previously documented in resin-embedded specimens [1, 18, 19] (B-D, white arrows), and, strikingly, thus far unreported tubular connections reaching up to ~300 nm (D-F, white arrowheads). Some DMVs were clearly interconnected through the ER (C, D) or by a tubular connection (H, black arrowheads). While these different connections were only detected in a subset of DMVs, it should be noted that only a percentage of an average-sized DMV (≤60%) would be contained in the thin cryo-lamellae used in this study (~150 nm thick). These observations are consistent with the previous notion of coronavirus-induced DMVs being largely interconnected as part of a reticulovesicular network [1, 18, 19]. (I) In some cases, DMVs appeared to fuse together through their outer membrane forming a so-called vesicle packet [18, 20]. (B-F, H, I) 7-nm thick tomographic slices. (G) 3D rendering of the tubular connection in (F).

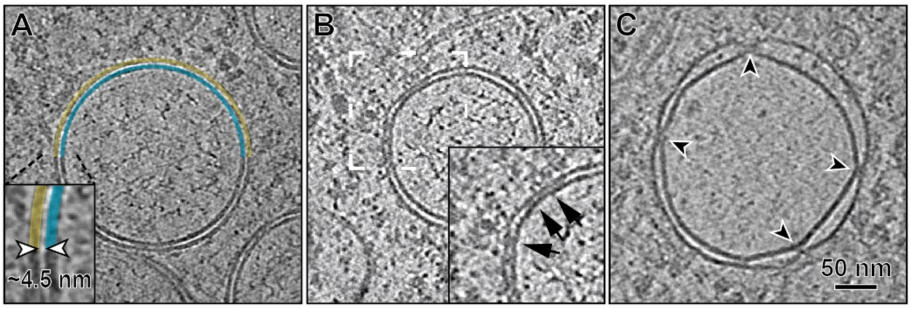


**Fig. S2.** Membrane-membrane interactions in coronaviral DMVs. (A) The outer and inner membranes of MHV-induced DMVs were most commonly separated by a regular spacing of ~4.5 nm. Such regular spacing is likely to be mediated by the interactions of factors embedded in the inner and outer membrane. (B) In fact, densities crossing the luminal space between the two membranes (boxed area, black arrows) were frequently found in these regions. The transmembrane proteins nsp3 and nsp4 have been previously implicated in membrane pairing [21-23]. Local deviations from the described regular inter-membrane spacing were common and, (C), occasionally, DMVs with largely dissociated membranes and molecular pores at constriction points (black arrowheads) were observed.

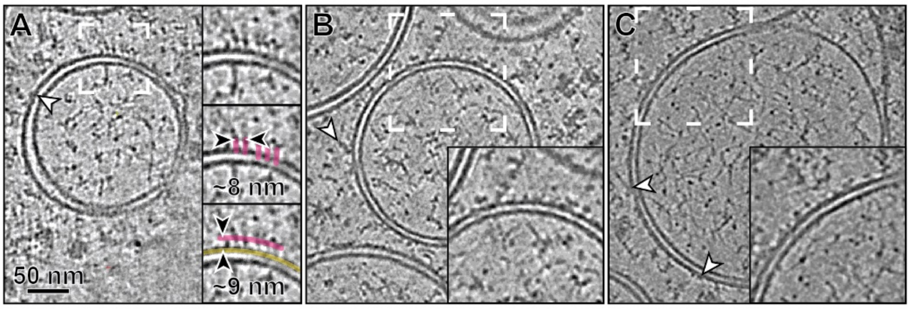


**Fig. S3.** Macromolecular features on the outer DMV membrane. (A-C) The outer DMV membrane appeared frequently decorated with a variety of molecular features (arrowheads and boxed areas). While the identity of these different densities cannot be ascertained, it is worth noting that a numerous viral and host proteins associate with coronavirus replication organelles [24] and that the large viral nsp3 transmembrane protein contains multiple cytosolic domains that have been implicated in interactions with host factors [25, 26]. A striking recurrent motif observed in the tomographic slices of the reconstructed DMVs were regular arrays (~8 nm spacing) of small globular masses, at ~9 nm distance from the outer membrane to which they seem to connect through a narrow link (boxed areas and insets).

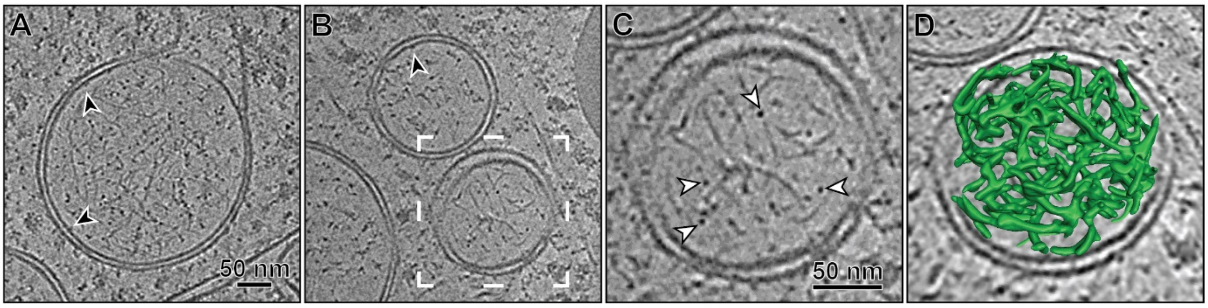


**Fig. S4.** DMV content. (A-C) DMVs appear to contain primarily filamentous structures, which can appear as dense points in the tomographic slices when viewed in cross-section (C, white arrowheads). The tomograms revealed relatively straight segments of a length consistent with the persistence length of dsRNA [27], which has been previously detected inside coronavirus-induced DMVs by immuno-EM [1, 18]. The dsRNA filaments seem to be loosely packed inside the DMVs forming what, at the current resolution, appears as an inextricable mesh of filaments. While ssRNA or protein complexes like the RTC could be contained in the DMV lumen, their presence within this intricate filamentous mesh could not be ascertained. The inner DMV membranes were relatively devoid of distinct associated partners, with only occasional small interacting densities (black arrowheads). (A-C) 7-nm thick tomographic slices. (D) 3D model of the filamentous content inside the DMV depicted in (C).

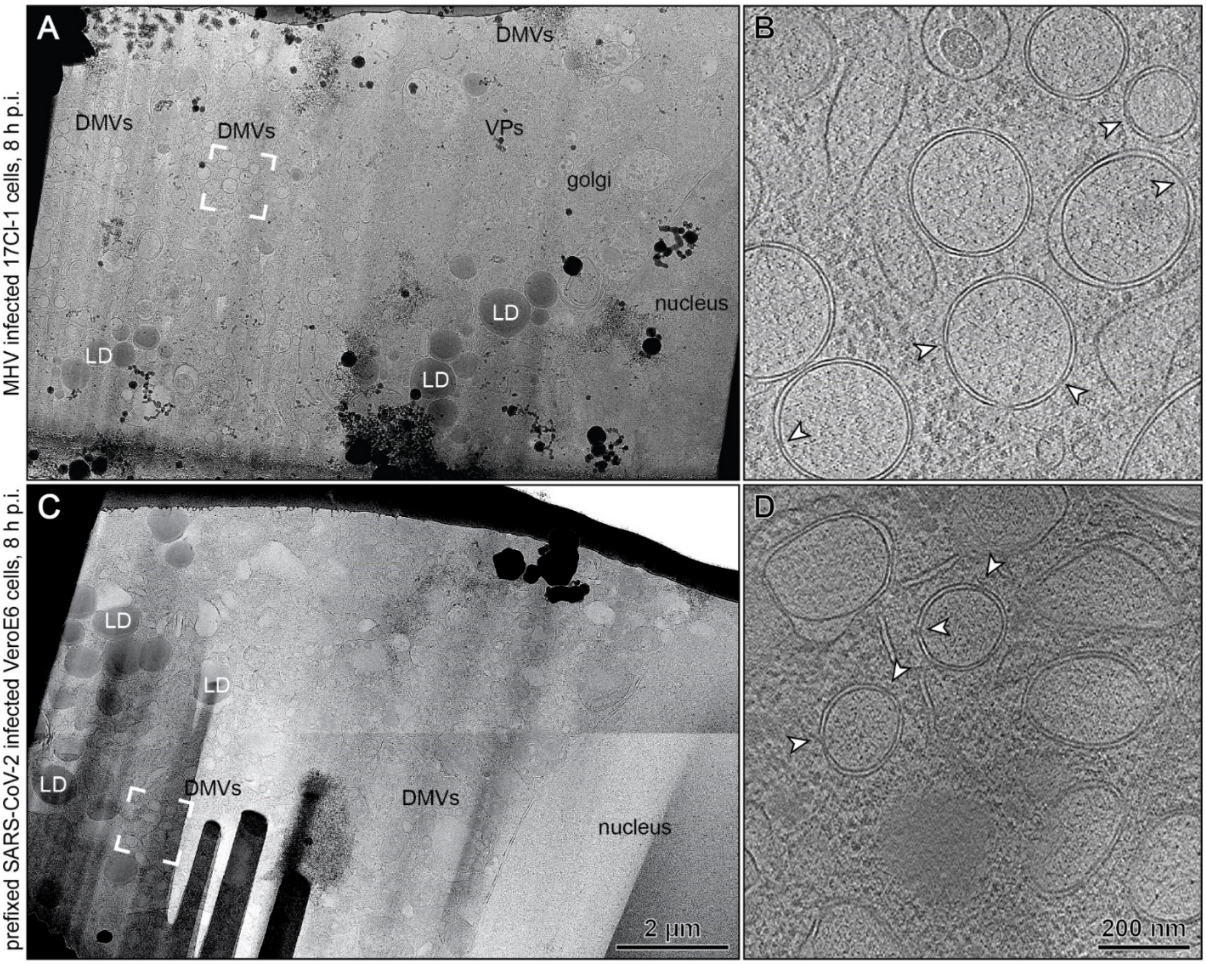


**Fig. S5.** Electron tomography on cryo-lamella of coronavirus-infected cells. Cryo-lamellae generated from cells infected with wt MHV (A, B) and cells infected with SARS-CoV-2 (C, D) that were chemically prefixed prior to vitrification. (A, C) Overviews of the cryo-lamellae, in which different structures, such as nucleus, mitochondria (M), the Golgi apparatus (Golgi), lipid droplets (LD), DMVs and virus particles (VPs), could be identified. (B, D) Slices (7 nm thick) of the tomograms reconstructed from tilt-series collected in the indicated regions (white frames in (A) and (C), respectively). Chemical fixation resulted in reduced contrast and produced morphological alterations in diverse organelles. Therefore, the partially distorted morphology of SARS-CoV-2-induced DMVs is likely also a result of fixation. Nonetheless, molecular pores in the DMV membranes could be identified for both samples (white arrowheads).

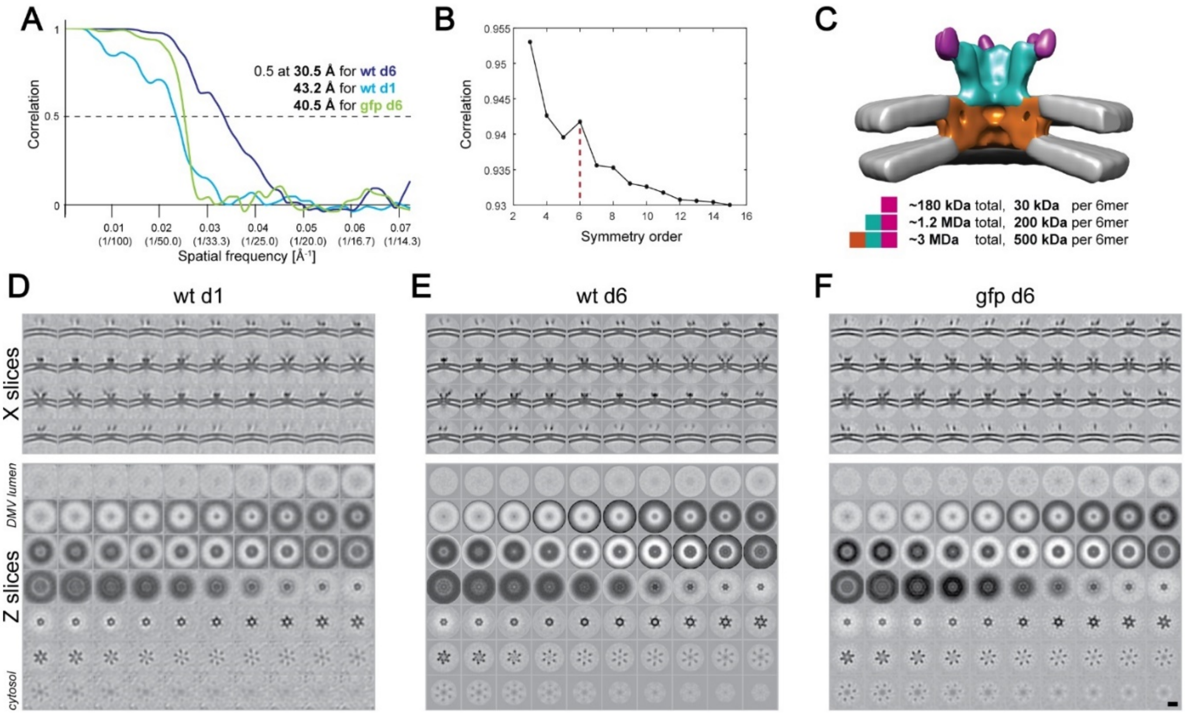


**Fig. S6.** Subtomogram averaging of the DMV molecular pore. (A) Fourier shell correlation curves from molecular pore particles aligned without imposing symmetry (d1) or implying 6-fold symmetry (d6). (B) A plot showing the correlation coefficients between unsymmetrized and symmetrized averages highlights the 6-fold symmetry of the pore complex. (C) Molecular masses of different sub-regions from the pore complex. (D-F) Slices (0.7 nm thick) throughout the molecular pore averages filtered to the achieved resolution, either parallel (top) or perpendicular (bottom) to the central axis of the complex. Wt, complex in wt MHV-induced DMVs; gfp, complex in MHV-Δ2-GFP-induced DMVs. Scalebar (D-F) 10 nm.

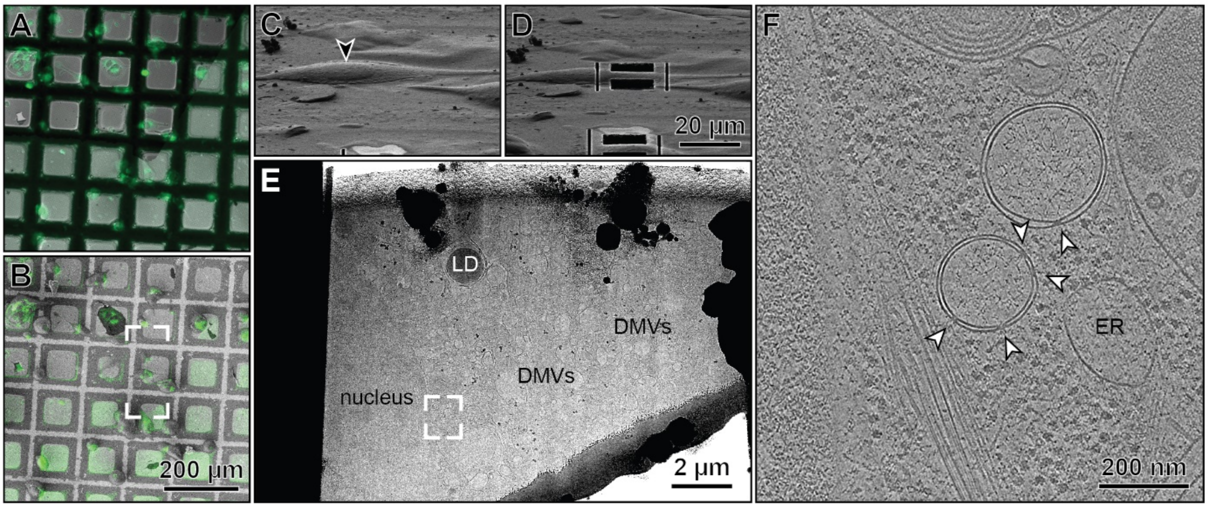


**Fig. S7.** Cryo-EM imaging of MHV-Δ2-GFP-infected cells was facilitated by cryo correlative light microscopy. (A) Fluorescence nsp3-EGFP signal was observed in perinuclear regions of plunge-frozen infected cells, which served as a maker for regions containing virus-induced replication compartments [18, 28]. (B) Overlays of the fluorescence signal on the corresponding SEM images guided the selection of suitable milling sites in the cryo-FIB-SEM machine. (C, D) The framed area in (B) as viewed from the FIB-beam direction before (C) and after (D) rough milling of the selected region. (E) Transmission EM overview of the thinned lamella, where regions of interest containing DMVs could be identified. The framed area was used for tilt-series acquisition. (F) A tomographic slice (7 nm thick) from the corresponding tomogram shows DMVs containing molecular pores (white arrowheads). LD, lipid droplet; ER, endoplasmic reticulum.

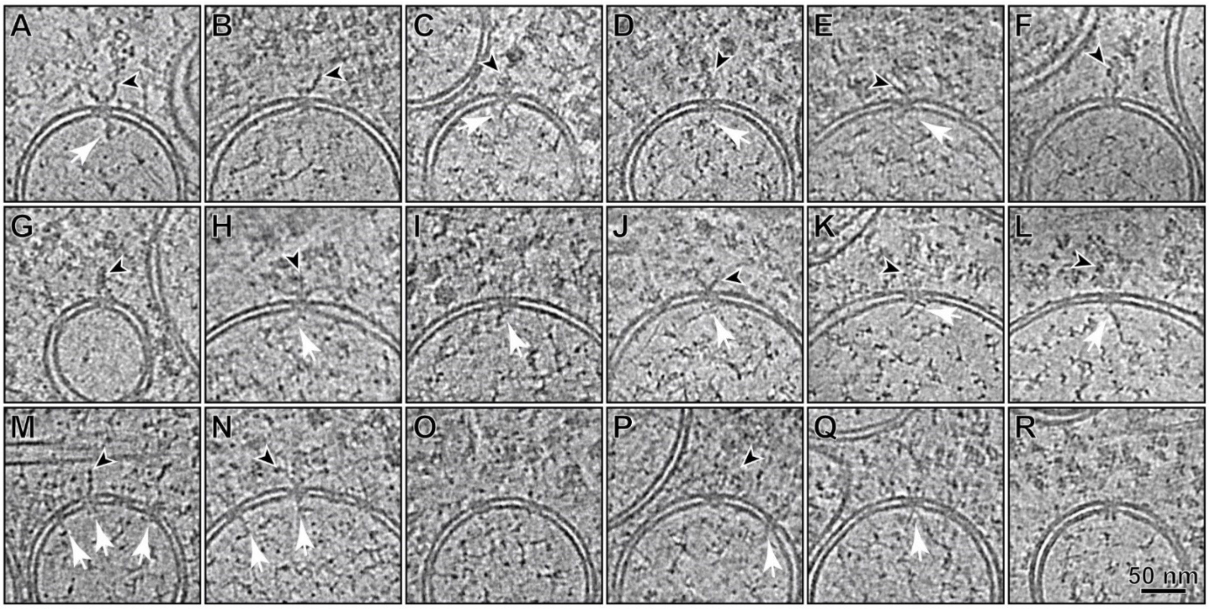


**Fig. S8.** Gallery of molecular pores from 7 nm thick tomographic slices. (A-Q) Various macromolecular structures appear to interact with both the cytosolic (top) and the DMV luminal (bottom) interfaces of the molecular pores. Cytosolic interaction partners (black arrowheads) range from (A, B, E) chain-like densities that seem to associate to the complexes prongs to larger assemblies that in some cases (D, G) are reminiscent of helical RNP. DMV luminal interaction partners (white arrows) appear to vary in mass and shape as well, while they frequently also show association to the DMV luminal filaments (C, D, I-L) assumed to be viral RNA. (R) In very few instances (less than 5 occurrences in a dataset containing more than 600 molecular pores) also pore complexes that appeared to be inverted in the DMV membrane have been observed.

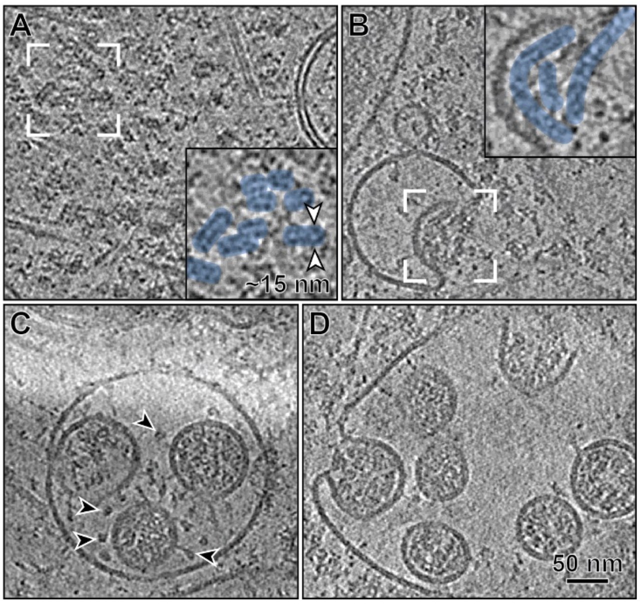


**Fig. S9.** RNP assemblies in the cytosol and the virions. Tomographic slices of 7 nm thickness showing (A) the abundant presence of cytosolic RNP assemblies in DMV-rich regions of the cytosol. The appearance of these structures is consistent with the helical RNPs of ~15 nm in diameter previously described by single-particle analysis [29], and with the RNP observed inside coronavirus particles in tomographic data (C, D) [30]. (B) RNP assemblies can associate to smooth-membrane compartments (boxed area) where they initiate the budding of virions into its the lumen [31]. (C-D) Budding virus particles containing RNP. Note also the spike protein on newly formed virions (C, black arrowheads).

**References:**


1. Snijder, E.J., et al., *A unifying structural and functional model of the coronavirus replication organelle: Tracking down RNA synthesis.* PLoS Biol, 2020. **18**(6): p. e3000715.
2. Freeman, M.C., et al., *Coronavirus replicase-reporter fusions provide quantitative analysis of replication and replication complex formation.* J Virol, 2014. **88**(10): p. 5319-27.
3. Graham, R.L., et al., *The nsp2 proteins of mouse hepatitis virus and SARS coronavirus are dispensable for viral replication.* Adv Exp Med Biol, 2006. **581**: p. 67-72.
4. Caly, L., et al., *Isolation and rapid sharing of the 2019 novel coronavirus (SARS-CoV-2) from the first patient diagnosed with COVID-19 in Australia.* Med J Aust, 2020. **212**(10): p. 459-462.
5. Wolff, G., et al., *Mind the gap: Micro-expansion joints drastically decrease the bending of FIB-milled cryo-lamellae.* J Struct Biol, 2019. **208**(3): p. 107389.
6. Hagen, W.J.H., W. Wan, and J.A.G. Briggs, *Implementation of a cryo-electron tomography tilt-scheme optimized for high resolution subtomogram averaging.* J Struct Biol, 2017. **197**(2): p. 191-198.
7. Zheng, S.Q., et al., *MotionCor2: anisotropic correction of beam-induced motion for improved cryo-electron microscopy.* Nat Methods, 2017. **14**(4): p. 331-332.
8. Rohou, A. and N. Grigorieff, *CTFFIND4: Fast and accurate defocus estimation from electron micrographs.* J Struct Biol, 2015. **192**(2): p. 216-21.
9. Xiong, Q., et al., *CTF determination and correction for low dose tomographic tilt series.* J Struct Biol, 2009. **168**(3): p. 378-87.
10. Frangakis, A.S. and R. Hegerl, *Noise reduction in electron tomographic reconstructions using nonlinear anisotropic diffusion.* J Struct Biol, 2001. **135**(3): p. 239-50.
11. Mastronarde, D., *Tomographic Reconstruction with the IMOD Software Package.* Microscopy and Microanalysis, 2006. **12**(S02): p. 178-179.
12. Castano-Diez, D., et al., *Dynamo: a flexible, user-friendly development tool for subtomogram averaging of cryo-EM data in high-performance computing environments.* J Struct Biol, 2012. **178**(2): p. 139-51.
13. Anger, A.M., et al., *Structures of the human and Drosophila 80S ribosome.* Nature, 2013. **497**(7447): p. 80-5.
14. Pettersen, E.F., et al., *UCSF Chimera--a visualization system for exploratory research and analysis.* J Comput Chem, 2004. **25**(13): p. 1605-12.
15. Harpaz, Y., M. Gerstein, and C. Chothia, *Volume changes on protein folding.* Structure, 1994. **2**(7): p. 641-9.
16. Zivanov, J., T. Nakane, and S.H.W. Scheres, *Estimation of high-order aberrations and anisotropic magnification from cryo-EM data sets in RELION-3.1.* IUCrJ, 2020. **7**(Pt 2): p. 253-267.
17. Lucic, V., A. Rigort, and W. Baumeister, *Cryo-electron tomography: the challenge of doing structural biology in situ.* J Cell Biol, 2013. **202**(3): p. 407-19.
18. Knoops, K., et al., *SARS-coronavirus replication is supported by a reticulovesicular network of modified endoplasmic reticulum.* PLoS Biol, 2008. **6**(9): p. e226.
19. Maier, H.J., et al., *Infectious bronchitis virus generates spherules from zippered endoplasmic reticulum membranes.* MBio, 2013. **4**(5): p. e00801-13.
20. Ogando, N.S., et al., *SARS-coronavirus-2 replication in Vero E6 cells: replication kinetics, rapid adaptation and cytopathology.* 2020: p. 2020.04.20.049924.

21. Angelini, M.M., et al., *Severe Acute Respiratory Syndrome Coronavirus Nonstructural Proteins 3, 4, and 6 Induce Double-Membrane Vesicles.* mBio, 2013. **4**(4): p. e00524-13.
22. Hagemeijer, M.C., et al., *Membrane rearrangements mediated by coronavirus nonstructural proteins 3 and 4.* Virology, 2014. **458-459**: p. 125-35.
23. Oudshoorn, D., et al., *Expression and Cleavage of Middle East Respiratory Syndrome Coronavirus nsp3-4 Polyprotein Induce the Formation of Double-Membrane Vesicles That Mimic Those Associated with Coronaviral RNA Replication.* mBio, 2017. **8**(6): p. e01658-17
24. V'Kovski, P., et al., *Determination of host proteins composing the microenvironment of coronavirus replicase complexes by proximity-labeling.* Elife, 2019. **8**: p. e42037.
25. Neuman, B.W., et al., *Proteomics analysis unravels the functional repertoire of coronavirus nonstructural protein 3.* J Virol, 2008. **82**(11): p. 5279-94.
26. Lei, J., Y. Kusov, and R. Hilgenfeld, *Nsp3 of coronaviruses: Structures and functions of a large multi-domain protein.* Antiviral Research, 2018. **149**: p. 58-74.
27. Abels, J.A., et al., *Single-molecule measurements of the persistence length of double-stranded RNA.* Biophys J, 2005. **88**(4): p. 2737-44.
28. Ulasli, M., et al., *Qualitative and quantitative ultrastructural analysis of the membrane rearrangements induced by coronavirus.* Cell Microbiol, 2010. **12**(6): p. 844-61.
29. Gui, M., et al., *Electron microscopy studies of the coronavirus ribonucleoprotein complex.* Protein Cell, 2017. **8**(3): p. 219-224.
30. Barcena, M., et al., *Cryo-electron tomography of mouse hepatitis virus: Insights into the structure of the coronavirion.* Proc Natl Acad Sci U S A, 2009. **106**(2): p. 582-7.
31. de Haan, C.A. and P.J. Rottier, *Molecular interactions in the assembly of coronaviruses.* Adv Virus Res, 2005. **64**: p. 165-230.
